# Assessment of microphytobenthos communities in the Kinzig catchment using photosynthesis-related traits, digital light microscopy and 18S-V9 amplicon sequencing

**DOI:** 10.1101/2024.06.17.599279

**Authors:** Ntambwe Albert Serge Mayombo, Mimoza Dani, Michael Kloster, Danijela Vidakovic, Dominik Buchner, Andrea M. Burfeid-Castellanos, Bánk Beszteri

**Affiliations:** Phycology Working Group, Faculty of Biology, University of Duisburg-Essen, Essen, Germany; University of Belgrade, Institute of Chemistry, Technology and Metallurgy, National Institute of the Republic of Serbia, Belgrade, Serbia; Aquatic Ecosystem Research, Faculty of Biology, University of Duisburg-Essen, Essen, Germany

**Keywords:** Microphytobenthos, biofilm, Bacillariophyta, environmental factors, digital microscopy, amplicon sequencing, photosynthetic biomass

## Abstract

Microalgae form an essential group of benthic organisms that respond swiftly to environmental changes. They are widely used as bioindicators of anthropogenic stressors in freshwater ecosystems. We aimed to assess the responses of microalgae communities to multiple environmental stressors in the Kinzig River catchment, home to a long-term ecological monitoring site, in Germany. We used a photosynthetic biomass proxy alongside community composition of diatoms assessed by digital light microscopy, and of microalgae by 18S-V9 amplicon sequencing, to characterise microalgae at 19 sampling sites scattered across the catchment. Our results revealed significant effects of physical and chemical factors on microalgae biomass and community compositions. We found that conductivity, water temperature and pH were the most important factors affecting microalgae community composition, as observed in both microscopy and amplicon analysis. In addition to these three variables, the effect of total phosphate on all microalgae, together with water discharge on the diatom (Bacillariophyta) communities, as assessed by amplicon analysis, may reveal taxon-specific variations in the ecological responses of different microalgal groups. Our results highlighted the complex relationship between various environmental variables and microalgae biomass and community composition. Further investigations, involving the collection of time series data, are required to fully understand the underlying biotic and abiotic parameters that influence these microalgae communities.

## 1 Introduction

The complex assemblages of microorganisms colonizing all submerged substrates also referred to as benthic microalgae biofilms, periphyton, or microphytobenthos (MPB), play an essential role in the biogeochemical cycles, primary production, and sediment stabilisation (B-Béres et al., 2023). At first glance, these tiny organisms may seem insignificant, but their presence and importance in rivers should not be underestimated. Through photosynthesis, these microscopic organisms act as microforests, converting sunlight into energy, and producing oxygen and organic matter. They contribute substantially to the global carbon and nitrogen cycles and provide essential ecosystem services that sustain diverse forms of life (Castro-Català et al., 2020; B-Béres et al., 2023). The environment and other organisms with which MPB biofilms interact (such as grazers) affect their species richness, community composition, and productivity (Birk et al., 2020). The ability of MPB biofilms to grow in different habitats and conditions also explains the compositional variability and complexity observed in their assemblages.

MPB assemblages are mainly dominated by diatoms (Bacillariophyta), but also other eukaryotes, including other microalgae such as Chlorophyta, Cyanobacteria and Euglenophyta, as well as other mixo-and heterotrophic protists, and fungi, may be present in lesser proportions (Underwood, 2010; Lemley et al., 2016; Dalu et al., 2020). Diatoms are secondary endosymbiotic eukaryotic algae belonging to the phylum Stramenopiles. Their glassy, siliceous cell walls known as frustules have been the focus of MPB community studies, mainly by observing their morphology under a light or a scanning electron microscope (Morales et al., 2001; Smol and Stoermer, 2010). Due to their short life cycles and rapid response to abiotic and biotic changes, MPB, particularly diatoms, are valuable bioindicators that can provide important insights into the health of aquatic ecosystems (Smol and Stoermer, 2010). Therefore, since 2000, the European Commission Water Framework Directive (European Commission, 2000) recommends the use of Biological Quality Elements (BQE), including diatoms, for the ecological assessment of surface waters.

Generally, studies of MPB biofilms have focused on the study of diatoms using light microscopy to analyse their community compositions. As diatoms are often the main component of these benthic systems, this approach has provided valuable insights into their ecological dynamics because they swiftly respond to both present and past environmental effects (Soininen et al., 2016; Viso and Blanco, 2023). Several studies have examined the responses of diatom communities to various anthropogenic disturbances (Walsh and Wepener, 2009; Hlúbiková et al., 2014; Teittinen et al., 2015; Stenger-Kovács et al., 2020). Diatom responses to a range of stressors, including heavy metal pollution (Lavoie et al., 2018), habitat fragmentation (Szczepocka et al., 2021) and climate change (Tornés et al., 2022) are well documented. This wealth of knowledge has greatly improved our understanding of diatom ecology and its resilience or vulnerability under different stressful conditions.

However, these traditional methods present a somewhat biased perspective. They often overlook the diversity and functional potential of the wider range of other organism groups that make up MPB biofilms, which includes not only other microalgae but also mixotrophic and heterotrophic protists and even fungi. This narrow focus on diatoms may lead us to underestimate the true ecological complexity and functional diversity of MPB biofilms. Additionally, it is not only the community composition that necessitates investigation but also ecosystem functions (Stenger-Kovács et al., 2020). Among these functions, photosynthesis stands out as a primary process underpinning the productivity and biogeochemical cycling in freshwater ecosystems.

A crucial aspect that remains underexplored is the potential differential impact of anthropogenic activities on microphytobenthos communities. As we continue to reshape the planet in the Anthropocene, understanding the nuanced effects of multiple anthropogenic stressors has become increasingly important. Human activities introduce different types of anthropogenic pressures in river catchments. These human-induced stressors have various effects on microphytobenthos communities (Feld et al., 2016; Birk et al., 2020). For example, in urban areas, with high population densities, extensive infrastructure and industrial activities, cocktail of point source pollutants may enter in freshwater ecosystems, that include wastewater treatment plant effluents, channel erosion, nutrients load, and inflow of contaminants such as heavy metals and other chemicals (Trábert et al., 2020).

This is in contrast with the rural landscapes dominated by agricultural land use, which is characterized typically by more diffuse pollution loads, such as the influx of nutrients, agrochemicals, pesticides and fine sediments, as well as hydromorphological alterations (Feld et al., 2016; Schürings et al., 2022). Yet, the effects of these multiple anthropogenic stressors (chemical and hydro-morphological) on microphytobenthos communities remain largely unknown.

In this study, we investigated how stream microphytobenthos communities respond to environmental variables in the Kinzig catchment, which is part of the Rhine-Main-Observatory (RMO), a long-term ecological research site (LTER) in central Germany. We focused on the primary environmental correlates of periphyton distribution in this river network. Our objectives were to: (1) assess the dynamics of photosynthetic biomass as a result of water physical and chemical conditions; (2) examine microalgae community composition and diversity metrics (abundances, richness, etc.) in relation to environmental variables and associated changes in ecological status; (3) identify the potential environmental factors that influence community composition in response to environmental conditions and ecological status, as compared to the macrozoobenthos studies of the LTER (Nguyen et al., 2023). We postulated that photosynthetic biomass and composition of MPB communities in the Kinzig catchment correlate with environmental factors. Higher biomass and reduced diversity will be associated with increased nutrient availability and eutrophication in the system. Similarly, hydrology (e.g. high flow velocity and depth, producing cloudy and low-light waters), will result also in a reduction in algal biomass and change in community composition. We seek to understand the resilience and adaptability of the MPB community by contrasting their responses to multiple anthropogenic stressors. In the face of numerous human pressures, this knowledge may help develop more potent conservation strategies for these essential freshwater ecosystem components. In addition to shedding light on the wider effects of multiple stressors on freshwater ecosystems, our study advances our understanding of MPB ecology.

## 2 Materials and methods

### 2.1 Study stie

In spring 2021, nineteen samples were collected from various streams across the Kinzig catchment and examined using a range of methods, including a recently developed digital version of diatom light microscopy for improved resolution of community composition (Burfeid-Castellanos et al., 2022), 18S-V9 amplicon sequencing for more comprehensive taxonomic characterisation of MPB communities (Amaral-Zettler et al., 2009) and in situ chlorophyll fluorescence measurements as a proxy for overall and group-specific photosynthetic biomass estimation (Kahlert and McKie, 2014).

The Kinzig catchment area (1058 km2) is a long-term ecosystem research site (LTER) in the Rhine-Main Observatory (RMO) (Nguyen et al., 2023). This low-mountain river system flows through different types of land use introducing various stressors. We are currently unaware of any ongoing restoration initiative in this river catchment, although land use in the study area has changed over time. Germany is actively working to reduce nutrient emissions in all catchment areas, in addition to ongoing efforts to address climate change (Nguyen et al., 2023). These factors could alter the structural composition of the microphytobenthos community over time. This study was part of the Collaborative Research Centre Multilevel Response to Stressor Increase and Release in Stream Ecosystems (CRC RESIST, https://sfb-resist.de/), which investigates the effects of global change in rivers and streams using field studies, mesocosm field and lab experiments and modelling approaches on several organism groups and scopes. We collected 19 samples in spring 2021 from ten tributaries selected within the Kinzig catchment (Figure 1). Sites were sampled based on adjacent land use.

**Figure 1.**
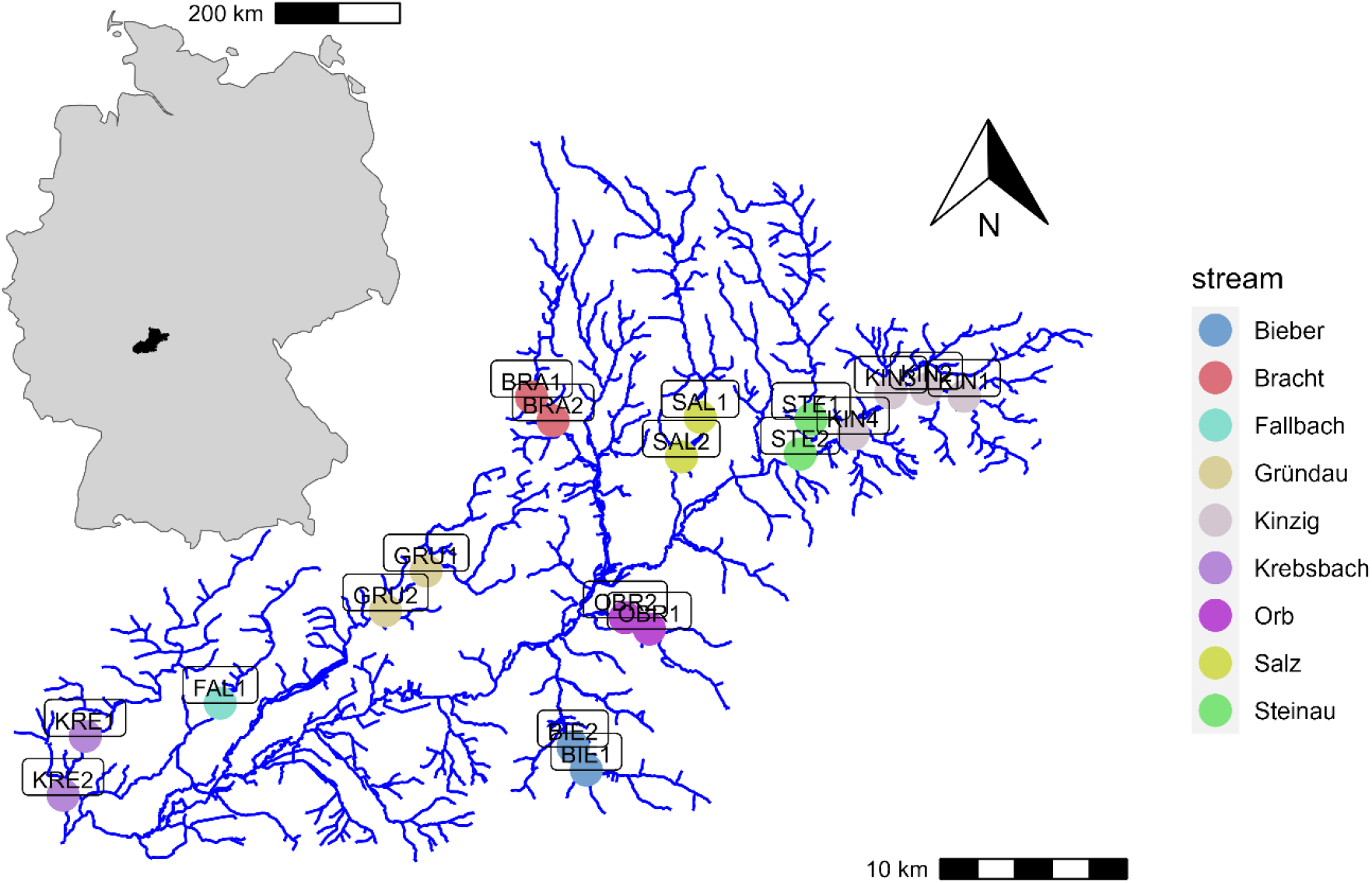
Map of the Kinzig catchment in Central Germany, showing sampling sites

### 2.2 Sample collection, preparation and processing

At least five separate chlorophyll (Chl) fluorescence measurements were made in situ on each sampled rock using a BenthoTorch (bbe Moldaenke GmbH, Schwentinental, Germany). The measurements were made using factory-programmed settings, and total Chl-a concentrations were given as the sum of biomass values for the three photosynthetic algal groups, namely cyanobacteria, green algae and diatoms. Biofilm samples were collected according to the recommendations and guidelines for sampling benthic diatoms from European rivers (Kelly et al., 1998; CEN. UNE-EN 13946:2014, 2014). A new, clean toothbrush was used to scrape approximately 20 cm2 from each of the five cobble stones sampled at each site. Approximately 20 mL of water from the river was used to collect the scrapings from the 5 cobble stones. As these samples were collected for both digital microscopy and DNA extraction, absolute ethanol (99%) was added, resulting in an approximate final concentration of approximately 75%, transported on ice and then frozen at -20°C once in the laboratory until they were divided for preparation for further analysis by digital microscopy and molecular methods. Water temperature, electrical conductivity and pH were measured in situ using a PCE-PHD 1 pH meter (Meschede, Germany). Other physical and chemical properties of the water including dissolved oxygen (DO), oxygen saturation, average velocity, average depth, discharge and water level were also measured in situ during biofilm sampling. In the laboratory, the Deutsches Institut für Normung (DIN) ISO norms were followed to determine the content of ortho-phosphate by photometry using a test kit, total phosphorus (TP) via ICP-MS (PerkinElmer), ammonia by photometry according to DIN 38406-5 (E5 – 1), nitrite by photozometry according to DIN EN 26 777, nitrate chloride and sulfate via ion chromatography, total nitrogen (TN) was calculate from nitrite, nitrate and ammonium values.

### 2.3 Diatom analysis by digital microscopy

Sample preparation and processing for digital microscopy followed the procedures described by Burfeid-Castellanos and coworkers (Burfeid-Castellanos et al., 2022). After five prewash cycles, the biofilm samples were digested with hydrogen peroxide and hydrochloric acid on a hot plate (Taylor et al., 2007). After seven rinsing cycles, centrifugation at 1200 rpm for 3 minutes (Eppendorf Centrifuge 5427 R; Eppendorf, Hamburg, Germany) and decanting and replenishing with distilled water, a 400 µL aliquot of each cleaned sample was pipetted onto a coverslip (15 x 15 mm, #1,5), allowed to dry and then mounted onto permanent glass slides using Naphrax resin (refractive index = 1.71, Biologie-Bedarf Thorns, Deggendorf, Germany). To produce virtual slides, an area of approximately 5 x 5 mm2 on the part of the permanent glass slide with the highest density of evenly distributed diatom valves was scanned using a VS200 slide scanner (EVIDENT, Tokyo, Japan) at a magnification of 600x (UPlanXApo) with the ASW 3.1 software (Olympus Soft Imaging Solutions GmbH, Münster, Germany).

These virtual slides were analysed for diatom counting and identification with the BIIGLE 2.0 web browser-based image annotation platform (Langenkämper et al., 2017; Zurowietz and Nattkemper, 2021) as described previously (Burfeid-Castellanos et al., 2022). Using a combination of general (Bey and Ector, 2013; Cantonati et al., 2017; Spaulding et al., 2022) and specific taxonomic literature (Cox, 1995; Levkov et al., 2013; Trobajo et al., 2013), at least 400 diatom valves were counted and identified at the highest level of possible taxonomic resolution, mostly at species level. For each sample, a list of the taxa and their relative abundances was generated to calculate the IPS (“Indice de Polluosensibilité Spécifique”) (Prygiel and Coste, 1993), using OMNIDIA software (Lecointe et al., 1993) and to run multivariate statistical analyses using R open source software environment for statistical analyses (R Core Team, 2023).

### 2.4 Molecular analyses

#### 2.4.1 DNA extraction and sequencing

Sample preparation and DNA extraction were performed following the silica bead-based extraction protocol described by Buchner (Buchner, 2022a; 2022b). Each sample was processed in 2 replicates. Genomic DNA amplification of the V9 hypervariable region of the small subunit ribosomal RNA genes (ca. 130 bp) was performed using the universal specific forward primer 1389F (5′-TTGTACACACCGCCC-3′) and the eukaryotic specific reverse primer 1510R (5′-CCTTCYGCAGGTTCACCTAC-3′) (Amaral-Zettler et al., 2009), following the protocol of Stoeck et al. (Stoeck et al., 2010). The resulting library was sequenced on an Illumina MiSeq at CeGat GmbH (Tübingen, Germany).

#### 2.4.2 Bioinformatics analysis

The Natrix2 workflow (Deep et al., 2023) was used to perform bioinformatics analyses on microphytobenthos Illumina amplicon sequencing data. The operational taxonomic units (OTUs) variant of the workflow was implemented using the SWARM v3.0.0 (Mahé et al., 2015) clustering algorithm. Paired-end reads were assembled using the simple Bayesian algorithm in PANDAseq v2.11 (Masella et al., 2012). Primers were trimmed with cutadapt v3.2 (Martin, 2011), and then filtered with a PANDAseq threshold score of 0.9, minimum length of 77, and maximum length 196 nucleotides. Sequence dereplication was performed using the CD-HIT v4.8.1 (Fu et al., 2012) algorithm at 100% similarity and chimeric sequences were identified and removed using VSEARCH v2.15.2 (Rognes et al., 2016). The split-sample approach (Lange et al., 2015) was applied to reduce erroneous sequences without strict abundance cut-offs. Statistical analysis of the amplicon sequencing data resulting from the split-sample process was conducted using the AmpliconDuo v1.1 R package (Lange et al., 2015). OTUs were generated by clustering sequences with Swarm v3.0.0 (Mahé et al., 2015). Then, mothur v1.40.5 (Schloss et al., 2009) was used to align OTUs against the protist ribosomal reference database (PR2) v.4.14.0 (Guillou et al., 2013). Finally, mumu (https://github.com/frederic-mahe/mumu), a C++ implementation of lulu (Frøslev et al., 2017), was used for post-clustering curation of the periphyton amplicon sequencing data. The replicates of sequenced samples were merged into one representative sample by calculating the sum of reads for each OTU and the negative controls with maximum reads were subtracted from the sum of both samples to standardize the proportion of sequences. This is important to reduce redundancy in the resulting dataset for subsequent statistical analyses. The resulting OTU table with their relative read abundances is used for subsequent multivariate statistical analyses.

### 2.5 Data analysis

We further examined the results from all above-described analyses using R v4.2.2 (R Core Team, 2023) and various packages including ’tidyverse’ v2.0.0 (Wickham et al., 2019), ’vegan’ v2.6-4 (Oksanen et al., 2022), ’factoextra’ v1.0.7 (Kassambara A, 2020), ’ggstatsplot’ v0.12.0 (Patil, 2021). Principal Components Analysis (PCA) was conducted after scaling the variables to have a mean of zero and a standard deviation of one, to show the correlation between environmental variables and sampling sites throughout the catchment. Correlation analysis was performed to determine the relationship between photosynthetic biomass and environmental variables, visualized using ’corrplot’ package (v0.92, Wei and Simko, 2021). We used the Hellinger transformed abundance matrix for species composition and OTU reads (Legendre and Gallagher, 2001). PERMANOVA was conducted to determine which environmental variables influenced diatoms and microalgae OTU assemblages. Furthermore, to characterise the influence of environmental variables on diatom species composition and microalgae OTU assemblages, we conducted the Redundancy Analysis (RDA). To simplify the RDA model, we performed a stepwise selection of statistically important environmental variables using ’ordiR2step()’ function. Only variables with significant correlations were visualized in the graphs.

## 3 Results

### 3.1. Environmental variables analysis

The physical and chemical parameters measured in this study are summarized in Table 1. The pH value of the water was neutral to relatively alkaline (7.03 to 9.55), with low to moderate electrical conductivity (108 to 862 µS/cm) (More details are given in Supplementary Table 1). Water temperature varied between 6 and 12.9°C, mean 8.69°C. The PCA performed on the matrix of physical and chemical variables showed that the first two axes explained 74.6% of the total variation in the data, so the first axis explained 58.2% of the variation and the second axis explained 16.4% of the variation (Figure 2). The most significant gradients in the environmental data were represented by chloride, conductivity, sulfate, nitrate and total nitrogen, which were the main contributors to the first PCA axis, while total phosphorus and average depth were the main contributor to the second PCA axis (Figure 2). The PCA ordination diagram showed two main groups of sampling locations. On the left side, the locations are distributed along a hydrological gradient (average depth vs. average velocity) that can be interpreted to roughly reflect altitude or upstream – downstream position in the catchment. The right side of the diagram shows three downstream sampling locations (FAL1, KRE1 and KRE2) offset from the others by increased nutrients and salinity (Figure 2, Table 1). This means that while our sampling locations seem to capture well the catchment-wide hydrological gradient, it does not resolve the nutrient and salinization gradient that appears close to orthogonal to the latter.

**Figure 2.**
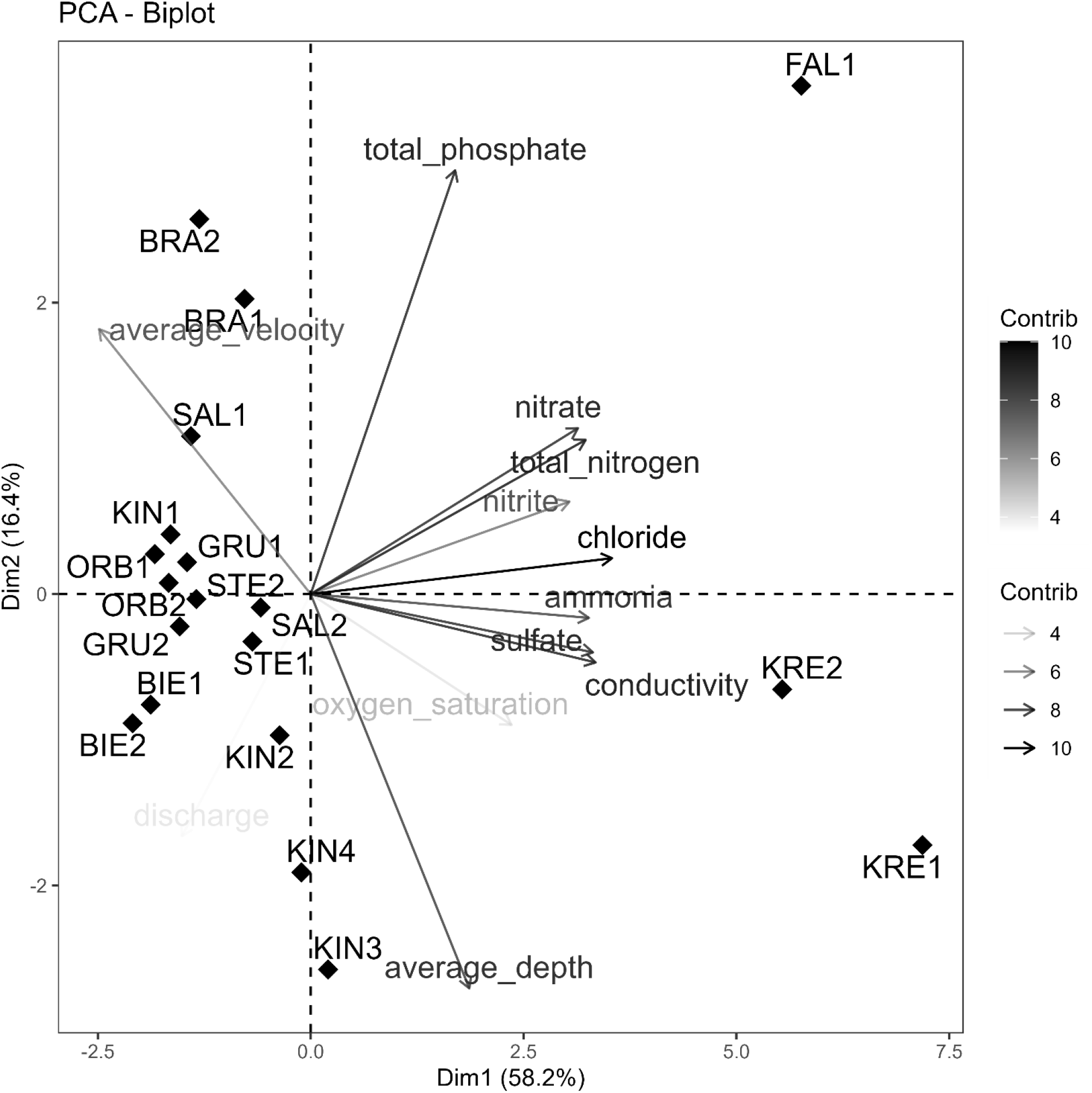
PCA biplot showing the relationship between the physical and chemical parameters and sampling sites The arrows and colour gradient show the contribution of different variables to the principal component.

**Table 1.**
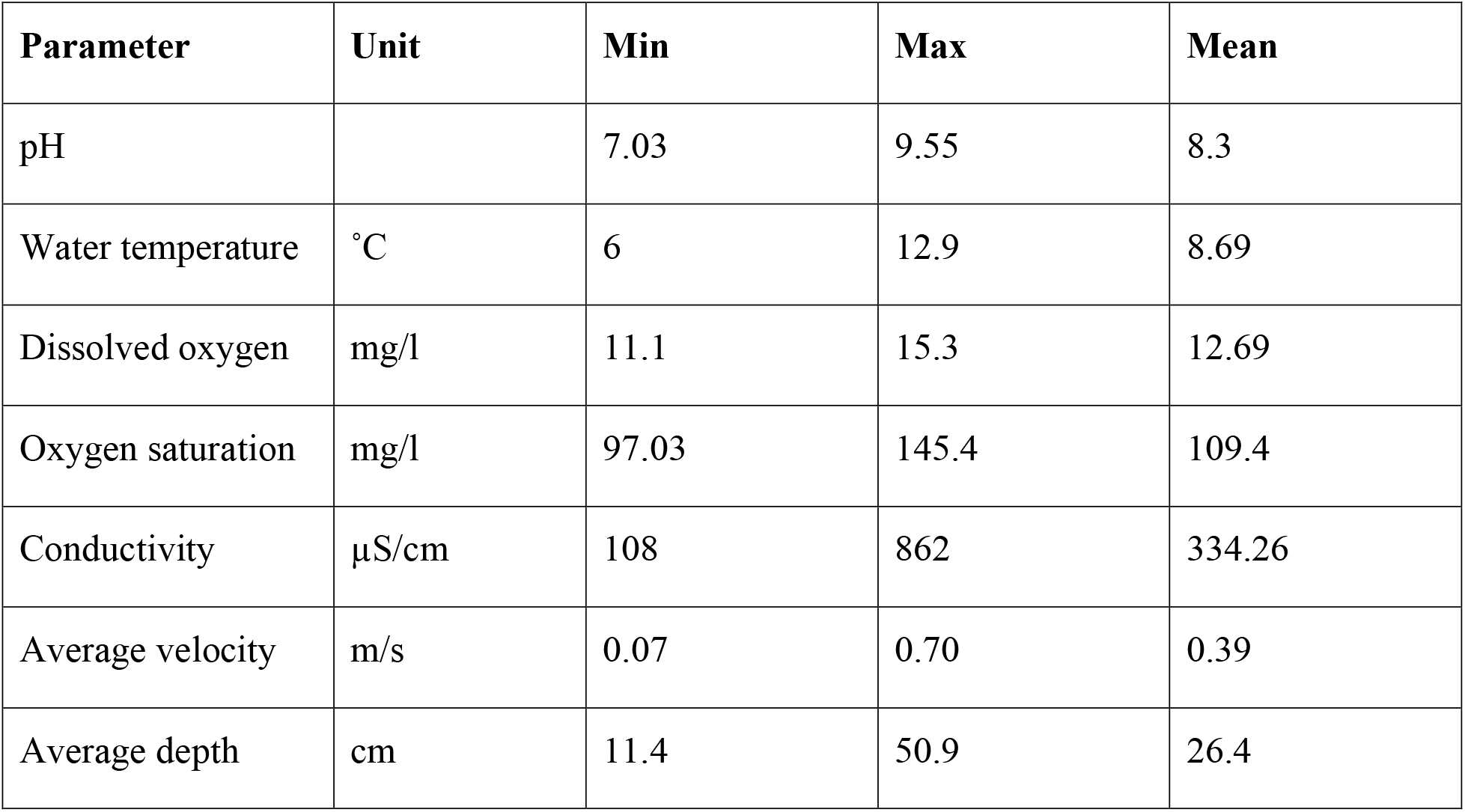
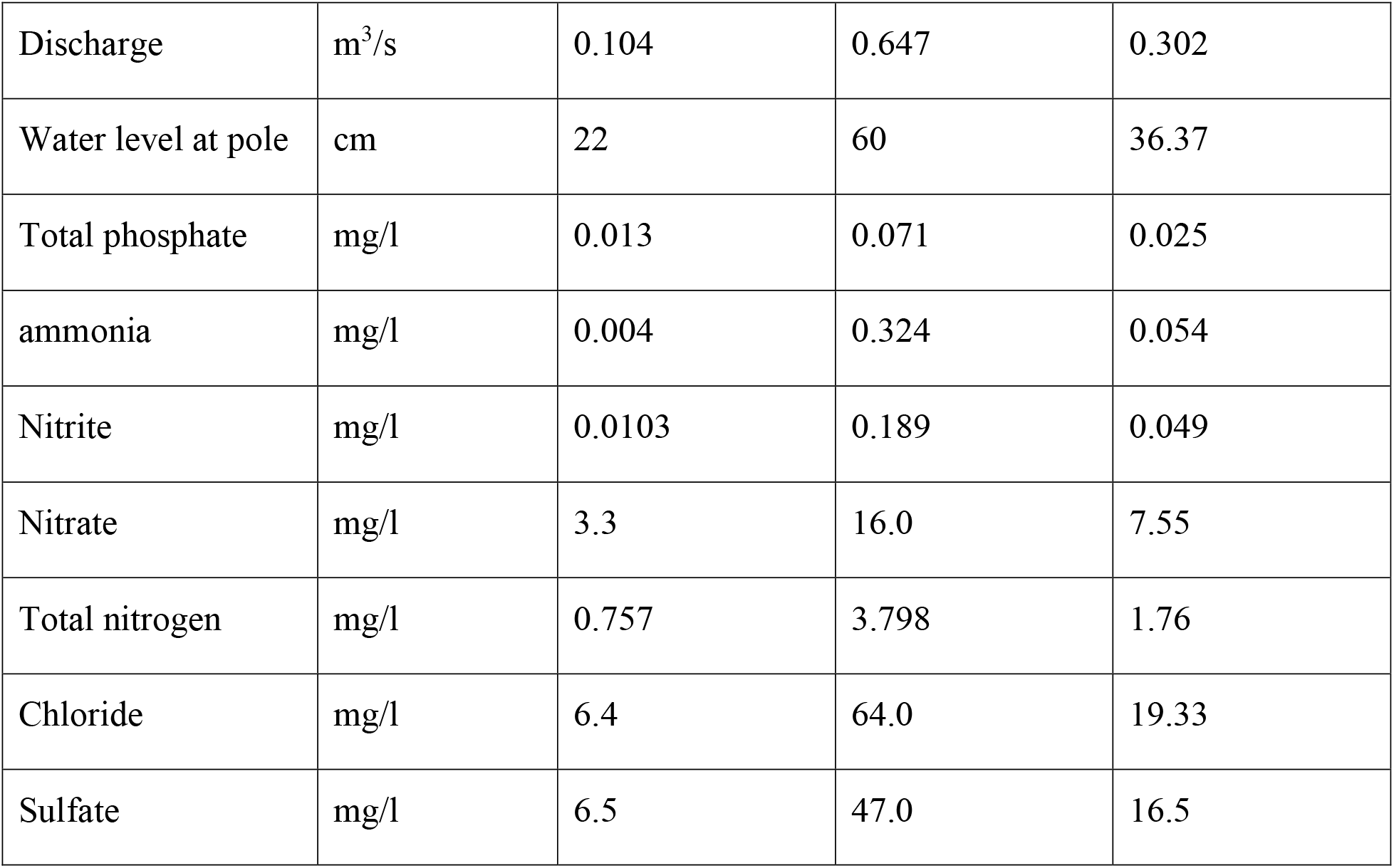
Summary statistics for physical and chemical parameters measured in the Kinzig catchment.

### 3.2. Photosynthetic biomass

We found statistically significant correlations between photosynthetic pigment fluorescence attributed by the BenthoTorch algorithm to cyanobacteria, green algae, diatoms and total algae concentration between the streams (ANOVA, all p values < 0.05). The correlation matrix of the physical and chemical environmental variables and mean photosynthetic biomass of different microalgae groups showed that pH was positively correlated with total and cyanobacterial fluorescence; sulfate strongly negatively correlated with total, cyanobactreial, and diatom (but not with green algal) chlorophyll signal. Total and green algal pigment fluorescence was also negatively correlated with average depth, as well as cyanobacterial and green algal signal with average velocity. Besides, water temperature and ammonia both showed a negative correlation with the diatom signal (Figure 3).

**Figure 3.**
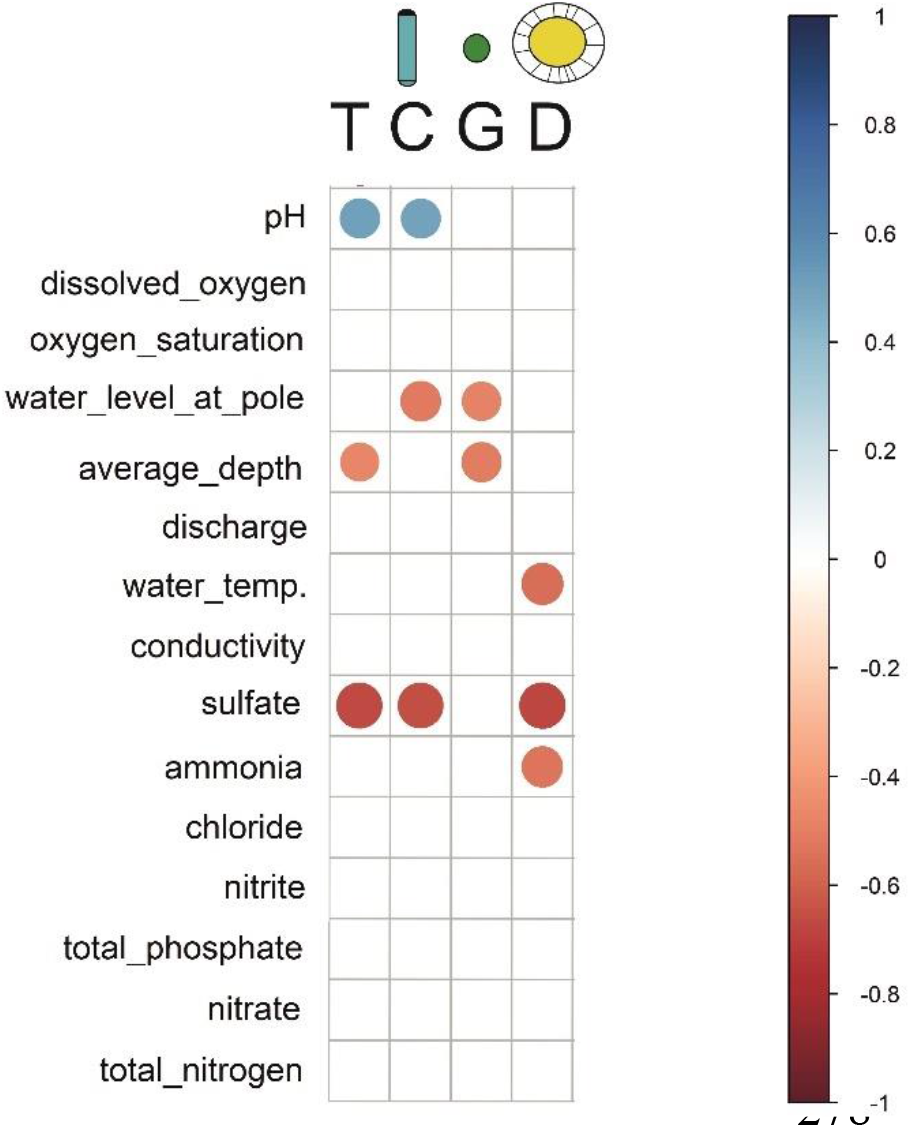
Correlation matrix showing the relationship between physical and chemical variables, as well as photosynthetic biomass of different microalgal groups. The variables in the matrix are ordered such that the correlated ones are next to each other. Positive correlations are displayed in blue and negative correlations in red colour. Colour intensity and the size of the circle are proportional to the correlation coefficients. Correlation coefficients with a p-value > 0.05 are considered insignificant and their fields remain blank. T = Total microalgae concentration, C = Cyanobacteria, G = Green algae, D = Diatoms

### 3.3. Diatom community composition: morphological analysis

A total of 142 diatom taxa belonging to 46 genera were identified by digital microscopy. Diatom community composition was dominated by *Navicula lanceolata*, *Navicula gregaria, Nitzschia dissipata, Achnanthidium minutissimum* and *Fragilaria deformis* (Figure 4 and Supplementary Table 2). Species richness of individual samples ranged from 21 to 56 (mean 37.3) and Shannon diversity index from 1.26 to 3.25 (mean 2.56). Both species richness and Shannon were lower in KRE1 and higher in GRU2. PERMANOVA common model including all predictors showed that conductivity, pH, and water temperature had significant effects (p < 0.05) on diatom community composition. All other factors had no statistically significant effects on diatom community composition (*p* > 0.05, Table 2).

**Figure 4.**
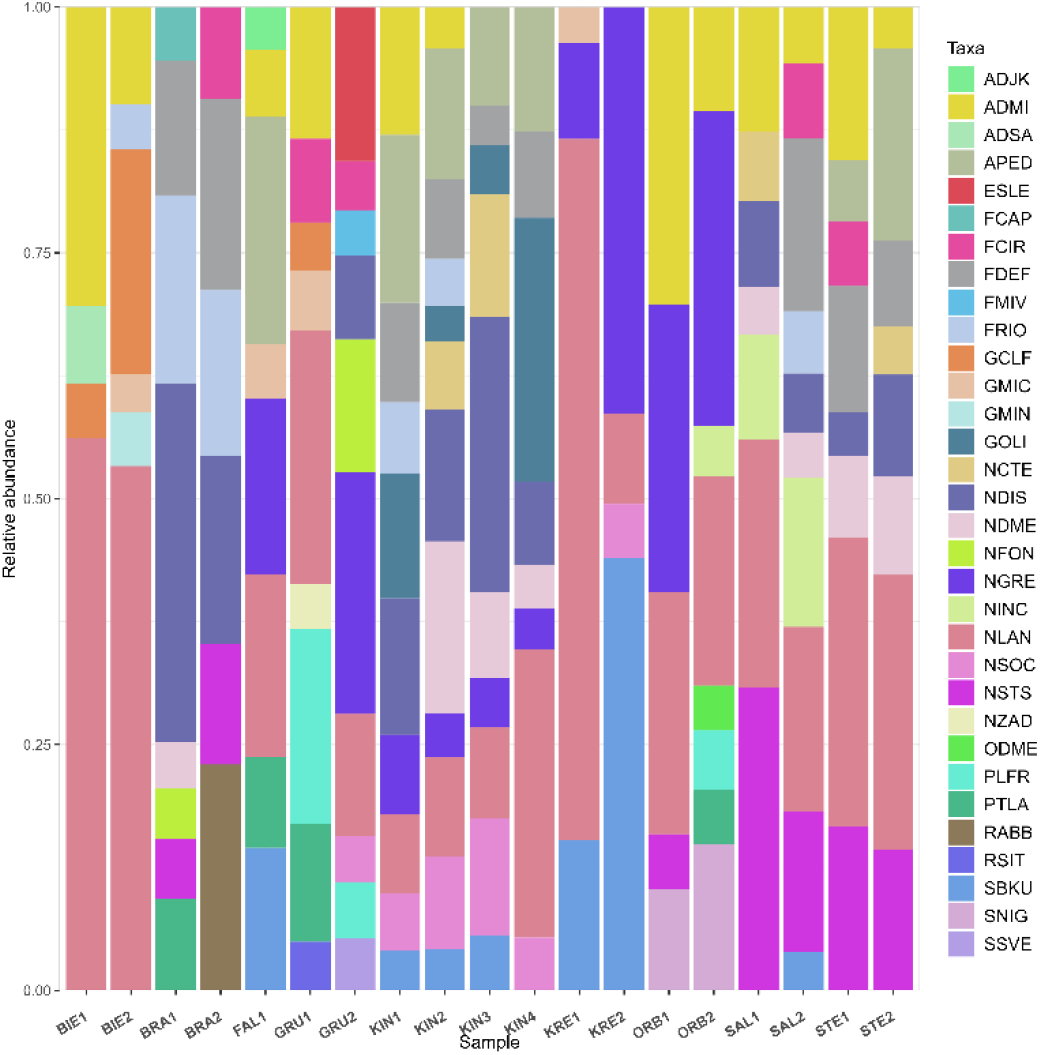
Relative abundance of dominant diatom taxa identified by digital light microscopy. Four letter code based on the OMNIDIA short code, explained in Supplementary Table 2.

**Table 2.** Results of PERMANOVA partitioning testing the effects of environmental variables on microalgae communities.

| Factor | df | SS | R<sup>2</sup> | F | P* |
| --- | --- | --- | --- | --- | --- |
| **Diatoms (digital microscopy)** |  |  |  |  |  |
| pH | 1 | 0.6901 | 0.1664 | 4.1280 | **0.007** |
| Water temperature | 1 | 0.6204 | 0.1496 | 3.7111 | **0.017** |
| Dissolved oxygen | 1 | 0.0741 | 0.0179 | 0.4431 | 0.860 |
| Oxygen saturation | 1 | 0.1741 | 0.0421 | 1.0445 | 0.416 |
| Conductivity | 1 | 0.5884 | 0.1419 | 3.5195 | **0.021** |
| Average velocity | 1 | 0.1455 | 0.0351 | 0.8700 | 0.535 |
| Average depth | 1 | 0.2064 | 0.0498 | 1.2345 | 0.311 |
| Discharge | 1 | 0.2811 | 0.0677 | 1.6816 | 0.195 |
| Water level at pole | 1 | 0.0822 | 0.0198 | 0.4920 | 0.808 |
| Total phosphate | 1 | 0.1957 | 0.0472 | 1.1706 | 0.355 |
| Ammonia | 1 | 0.1018 | 0.0245 | 0.6086 | 0.772 |
| Nitrite | 1 | 0.0713 | 0.0172 | 0.4263 | 0.863 |
| Nitrate | 1 | 0.1581 | 0.0381 | 0.9457 | 0.484 |
| Chloride | 1 | 0.0840 | 0.0202 | 0.5022 | 0.801 |
| Sulfate | 1 | 0.1728 | 0.0417 | 1.0334 | 0.437 |
| Residual | 3 | 0.5016 | 0.1209 |  |  |
| Total | 18 | 4.1481 | 1.0000 |  |  |
| **Diatom OTUs (18S-V9 amplicon sequencing)** |  |  |  |  |  |
| pH | 1 | 0.2665 | 0.1329 | 5.0575 | **0.001** |
| Water temperature | 1 | 0.2264 | 0.1129 | 4.2969 | **0.001** |
| Dissolved oxygen | 1 | 0.0701 | 0.0350 | 1.3306 | 0.267 |
| Oxygen saturation | 1 | 0.0927 | 0.0462 | 1.7598 | 0.105 |
| Conductivity | 1 | 0.3972 | 0.1981 | 7.5385 | **0.001** |
| Average velocity | 1 | 0.0465 | 0.0232 | 0.8822 | 0.563 |
| Average depth | 1 | 0.1051 | 0.0524 | 1.9947 | 0.069 |
| Discharge | 1 | 0.1335 | 0.0666 | 2.5337 | **0.018** |
| Water level at pole | 1 | 0.0477 | 0.0238 | 0.9044 | 0.505 |
| Total phosphate | 1 | 0.1350 | 0.0674 | 2.5629 | **0.021** |
| Ammonia | 1 | 0.0788 | 0.0393 | 1.4956 | 0.190 |
| Nitrite | 1 | 0.0474 | 0.0236 | 0.8989 | 0.513 |
| Nitrate | 1 | 0.0894 | 0.0446 | 1.6974 | 0.112 |
| Chloride | 1 | 0.0651 | 0.0325 | 1.2359 | 0.290 |
| Sulfate | 1 | 0.0457 | 0.0228 | 0.8665 | 0.552 |
| Residual | 3 | 0.1581 | 0.0788 |  |  |
| Total | 18 | 2.0050 | 1.0000 |  |  |
| **All microalgae OTUs (18S-V9 amplicon sequencing)** |  |  |  |  |  |
| pH | 1 | 0.4197 | 0.1389 | 4.6839 | **0.001** |
| Water temperature | 1 | 0.2914 | 0.0964 | 3.2521 | **0.005** |
| Dissolved oxygen | 1 | 0.1750 | 0.0579 | 19529 | 0.052 |
| Oxygen saturation | 1 | 0.1454 | 0.0481 | 1.6226 | 0.116 |
| Conductivity | 1 | 0.5216 | 0.1726 | 5.8201 | **0.001** |
| Average velocity | 1 | 0.0914 | 0.0303 | 1.0205 | 0.449 |
| Average depth | 1 | 0.1524 | 0.0504 | 1.7011 | 0.079 |
| Discharge | 1 | 0.1526 | 0.0505 | 1.7028 | 0.080 |
| Water level at pole | 1 | 0.0947 | 0.0313 | 1.0569 | 0.419 |
| Total phosphate | 1 | 0.1782 | 0.0590 | 1.9885 | **0.044** |
| Ammonia | 1 | 0.1373 | 0.0454 | 1.5326 | 0.116 |
| Nitrite | 1 | 0.0822 | 0.0272 | 0.9171 | 0.564 |
| Nitrate | 1 | 0.1209 | 0.0400 | 1.3489 | 0.191 |
| Chloride | 1 | 0.1103 | 0.0365 | 1.2310 | 0.283 |
| Sulfate | 1 | 0.0800 | 0.0265 | 0.8927 | 0.568 |
| Residual | 3 | 0.2688 | 0.0890 |  |  |
| Total | 18 | 3.0217 | 1.0000 |  |  |

*df*: degrees of freedom; *SS*: sum of squares; $R^2$ : variance explained by the groups, *F*: *F*-statistic, *p*: p- values, \*significant p-values are marked in bold

The RDA model explained a variation of 27% on the first axis and 17.8% on the second. The environmental factors pointed out by this analysis to show the most statistically significant association (p < 0.05) with the differentiation of diatom communities were pH, water temperature, oxygen saturation (oxygen_100%), conductivity, average velocity, average depth, ammonia, chloride and sulfate. Of these, only pH, water temperature and conductivity showed a strong effect, while the effects of all the other factors was weak. The RDA ordination diagram shows sites in the upper left quadrant that are largely influenced by pH and oxygen saturation, and were associated with *Nitzschia soratensis* and *Fragilaria rinoi*. Sites plotted in the lower right quadrant, were mainly influenced by ammonia, sulfate, chloride, conductivity and average depth. These sites were associated with taxa such as *Surirella brebissonii* var. *kuetzingii* and *Navicula gregaria* (Figure 5). The ecological status index of diatoms indicated by the IPS ranged from 10.5 to 15.9 (mean 14.3), indicating poor to good ecological status. Only one site, KRE2 had poor ecological status (Figure 5). The main component of the environmental gradient correlated with community composition was that of water temperature vs. pH and oxygen saturation, while the second axis was most strongly correlated with water depth, salinity components (chloride, sulfate and conductivity), and ammonia concentration. Interestingly, the magnitude of differentiation in diatom community composition appeared lower than the separation in environmental variables between the three high-nutrient, high-sulfate locations (FAL1, KRE1, KRE2) observed in the above PCA.

**Figure 5.**
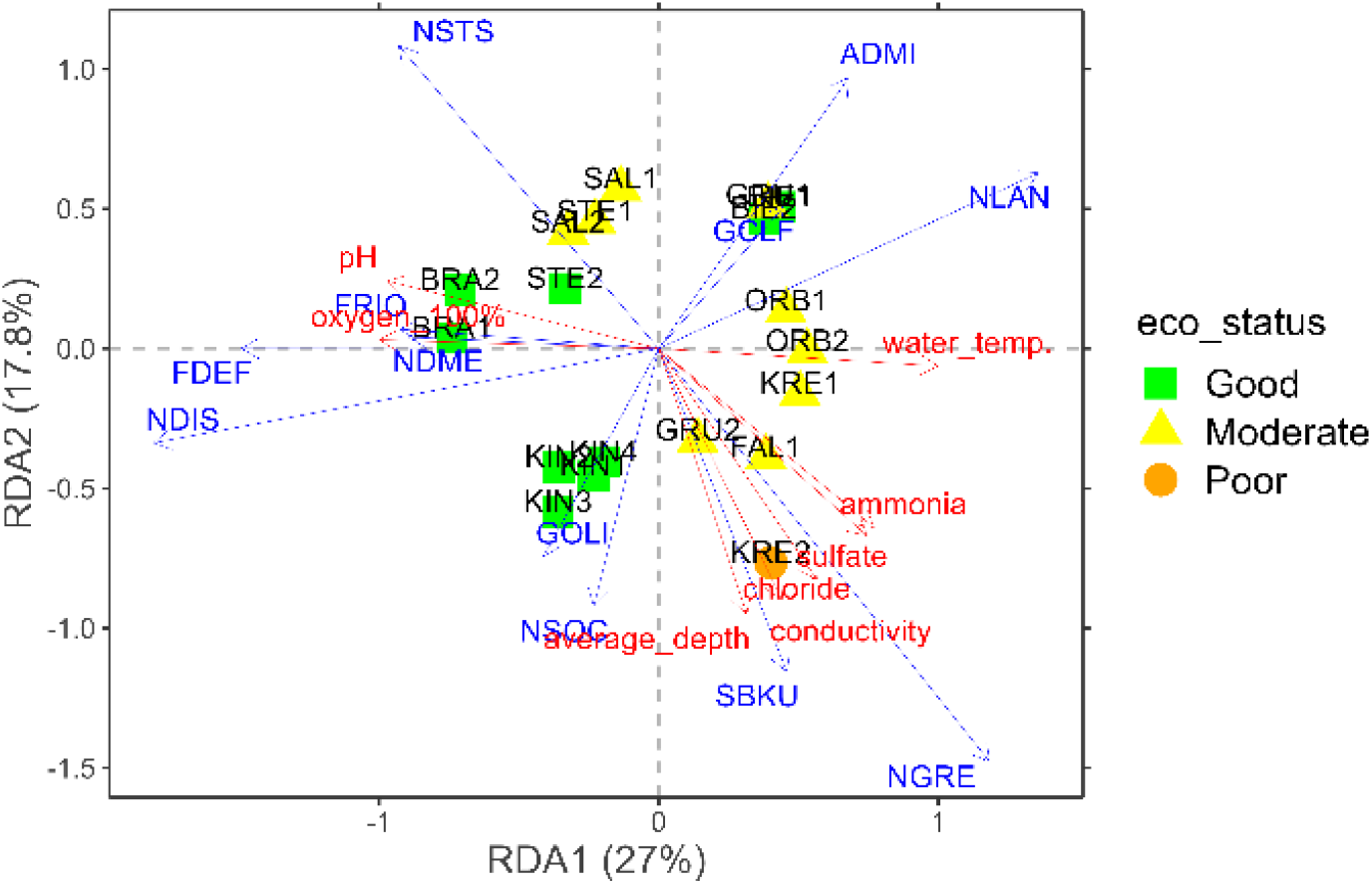
Redundancy analysis (RDA) plot presenting the relationships between the analysed diatom assemblages in the samples based on digital light microscopy and environmental variables. Four letter code based on the OMNIDIA short code, explained in Supplementary Table 2.

### 3.4. Microalgae community composition: 18S-V9 amplicon sequencing

A total of 21,293,230 reads were obtained after running the Natrix Bioinformatics pipeline on 18S-V9 amplicon sequencing data from the 19 sampling sites (average 1,120,696 reads per sample, minimum 533,819 reads and maximum 1,604,630 reads). These reads were clustered into 7294 OTUs from various protist groups and assigned taxonomy up to the highest possible level. Microalgae OTU richness ranged from 803 to 2295 (mean 1460), Shannon diversity from 6.69 to 7.74 (mean 7.24). The lowest alpha diversity values were recovered in STE1 and the highest in GRU2. Diatoms contributed most of the MPB OTUs, making up the majority of reads (13,322,277 or about 79.3% of total read counts; Figure 6A, Supplementary Table 2). We found 1400 different diatom OTUs, with OTU richness ranging from 140 to 392 (mean 247.59), Shannon diversity ranging from 1.52 to 2.66 (mean 2.16). The lowest value of diatom OTU richness was recorded in BIE2 and the highest in FAL1. The diatom OTUs with the highest read counts were assigned to *Navicula radiosa*, *Navicula gregaria*, *Nitzschia dissipata*, *Surirella brebissoni*, *Melosira varians* and *Asterionella formosa* (Figure 6B). Many of these taxa were also observed during the digital light microscopy analysis, except for *N. radiosa* and *A. formosa* (Supplementary Table 2).

**Figure 6.**
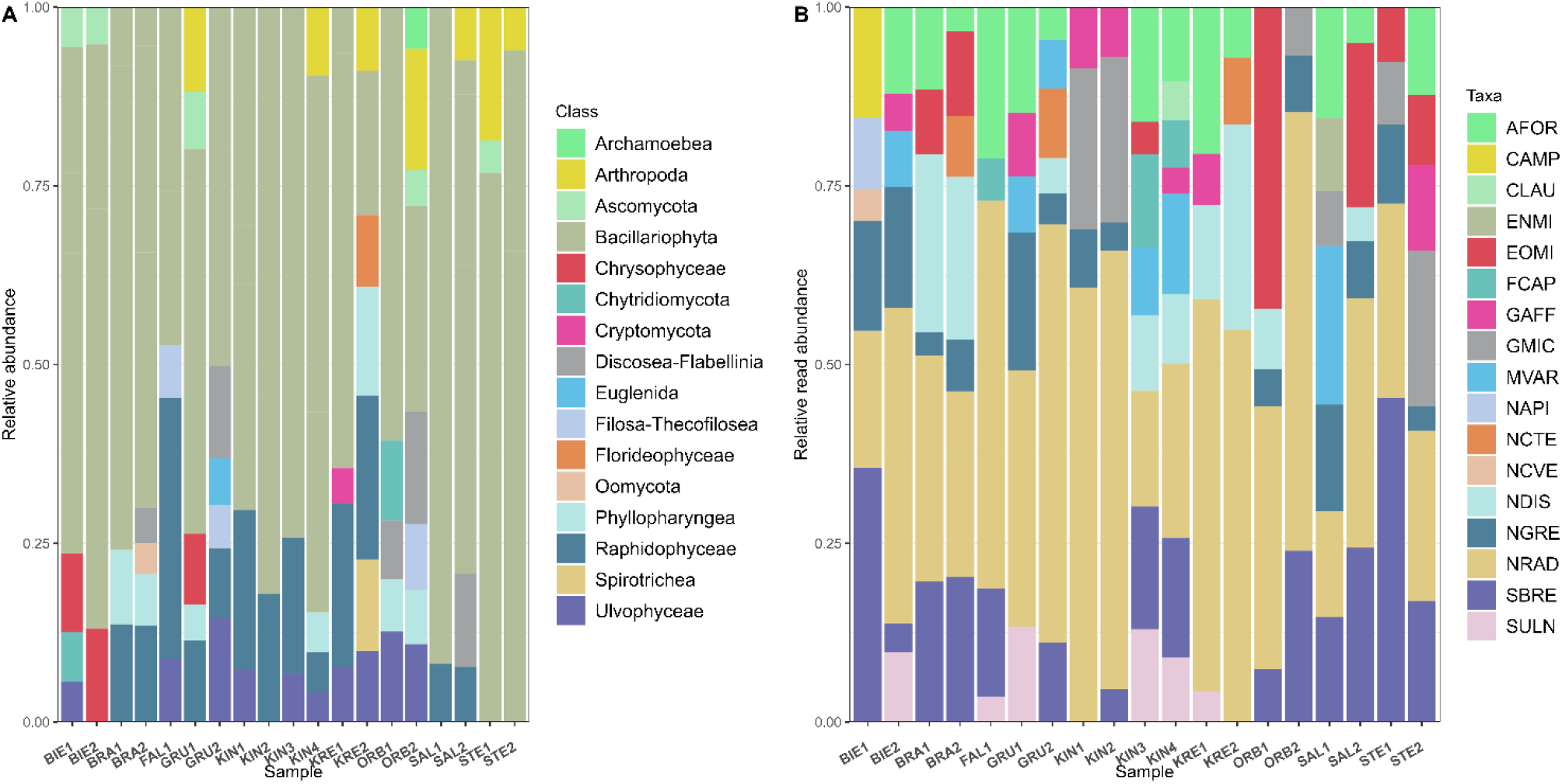
Relative reads abundance of **(A)** dominant OTU classes in the samples and **(B)** taxa associated with dominant diatom (Bacillariophyta) OTUs. Four letter code based on the OMNIDIA short code, explained in Supplementary Table 2.

PERMANOVA common model including all predictors run on the full microalgae OTU dataset revealed that water pH, temperature, conductivity had significant effect on community composition (*p* < 0.05, Table 2). Total phosphate had a weak effect on microalgae community. Similarly, the same model run on the diatom subset of OTU reads revealed significant effects of water pH, temperature, conductivity, discharge and total phosphate (*p* < 0.05, Table 2).

The RDA model examining the relationship between the diatom subset of OTU reads and environmental variables showed a lower explanatory power than the one for the diatom microscopy data set, with a fitted variation of 20.5% explained by the first axis and 17% by the second axis (Fig 7A). The major explanatory gradient here was a salinization – depth gradient that played a slightly lower role in explaining the diatom microscopy results, with the pH vs. water temperature gradient occurring here along the second axis (Fig 7A). In the upper left quadrant of the RDA ordination diagram, sites such as OBR1 and OBR2 were influenced by flow velocity, while pH was the most important predictor at sites such as KIN2 and KIN3 in the lower left quadrant. Sites BIE1, BIE2 and GRU1 in the upper right quadrant were mostly associated with higher water temperature. Finally, FAL1, KIN1 and KIN2 in the lower right quadrant were mainly influenced by chloride, conductivity and depth. The ecological status index (IPS) calculated from the diatom amplicon data set was between 11.4 and 18.1 (mean value 15.6), indicating poor to very good water quality (one has to note that this index was not calibrated for amplicon data sets). Only site KRE2 had a poor ecological status (Figure 7A). In this RDA, the high nutrient, salinized set of samples (FAL1, KRE1, KRE2) appeared with the highest RDA1 scores, but not so strongly offset from the rest of the samples as in the environmental PCA.

**Figure 7.**
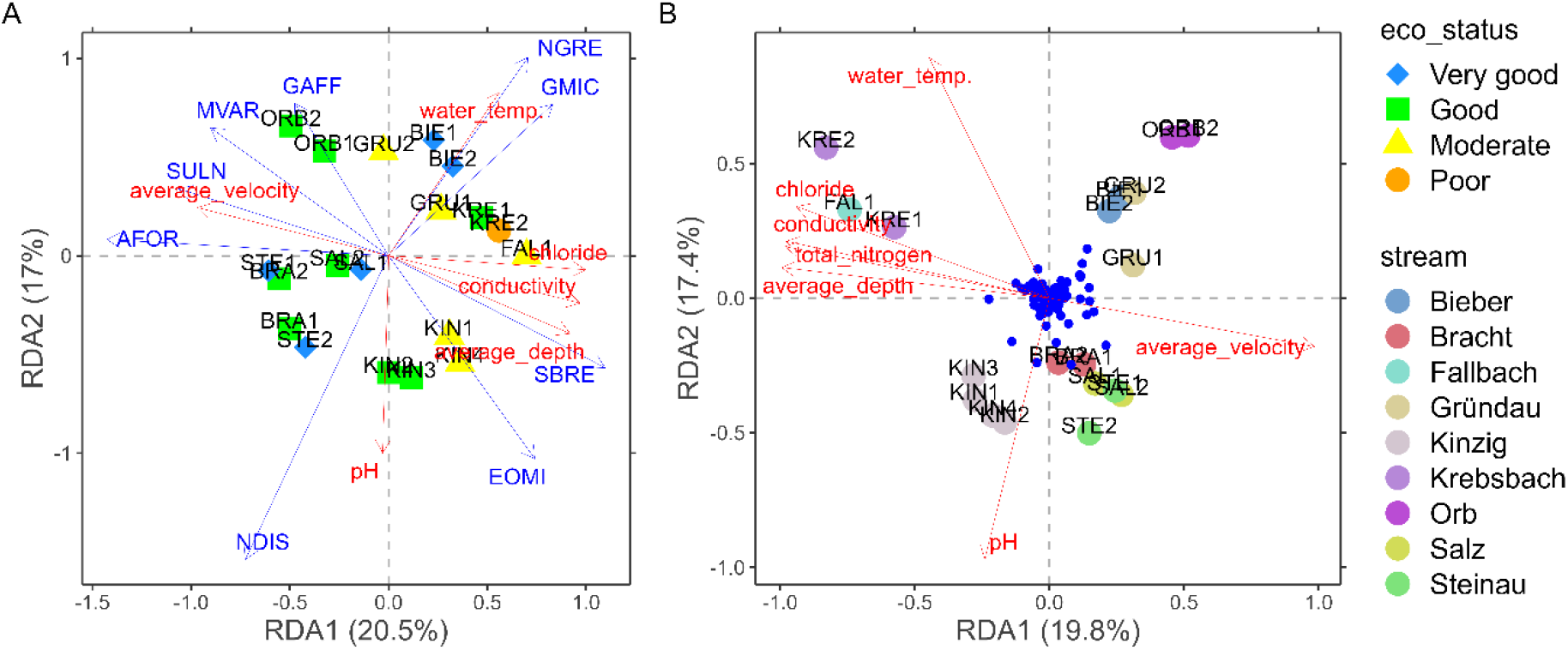
Redundancy analysis (RDA) plots presenting the relationships between the analysed OTUs assemblages and environmental variables: **(A)** diatom (Bacillariophyta) subset of OTUs (ecological status according to IPS index) and **(B)** the whole microalgae OTU dataset (small blue dots show significant OTUs). Four letter code based on the OMNIDIA short code, explained in Supplementary Table 2.

Similarly, the RDA model explained a fitted variation of 19.8% on the first axis and 17.4% on the second axis when analysing the relationship between the entire microalgae OTU read dataset and physical and chemical variables (Figure 7B). The main explanatory environmental gradients were highly similar to the diatom OTU data set (with opposite signs along the first axis). Major environmental factors contributing to these variations (p < 0.05) included pH, temperature, conductivity, average velocity, average depth, total nitrogen and chloride. The separation of FAL1, KRE1 and KRE2 from the rest of the samples was clearer in this case than for both above diatom community data sets. (Figure 7B). Sites located in the Kinzig Stream (KIN1 – KIN4), plotted in the lower left quadrant, were mainly influenced by pH, while sites plotted in the lower right quadrant (BRA1, BRA2, SAL1, SAL2, STE1 and STE2) were associated with flow velocity (Figure7B).

## 4 Discussion

### 4.1 Microalgae community composition in response to environmental variables

The effort to understand the effects of anthropogenic stressors on microalgal communities within a river catchment remains at the forefront of research in aquatic ecosystems. The compelling reason for this increasing interest is the widespread recognition that anthropogenic activities may have differential impacts on freshwater ecosystems. This study aimed to identify the main environmental drivers of variability in photosynthetic biomass and microalgal community compositions in the anthropogenically stressed Kinzig catchment. Alongside a set of candidate environmental predictors, we used for this purpose data sets capturing total and group-specific photosynthetic biomass and community composition of diatoms (characterized by microscopy vs. by amplicon sequencing) as well as of all microalgal groups (amplicon sequencing).

The PCA ordination of environmental predictors highlighted a hydrological gradient (depth and discharge vs. velocity) well resolved by our samples. Close to orthogonal to that, the PCA indicated a water chemistry gradient (nutrient and salts, the latter mainly sulphate, but also chloride) captured by two clusters of sites (the high nutrient, high salinity locations FAL1, KRE1 and KRE2 vs. all others). In addition, total phosphate concentration showed an aspect of variability somewhat decoupled from both of these gradients (Fig 2).

In terms of photosynthetic biomass (as estimated using a BenthoTorch), the depth gradient was slightly, but significantly negatively associated with total and green algal chlorophyll concentration (Fig. 3.), indicating a slightly decreasing trend in green algal biomass content with increasing depth which could be due to light limitations (Virtanen and Soininen, 2012). Total, cyanobacterial and diatom chlorophyll signals were strongly negatively correlated with sulfate, also being the main driver of the salinity gradient in the system. Thus, whereas biomass of green algae seems to respond more strongly along the hydrological gradient, complementary to this, the chlorophyll biomass of diatoms and cyanobacteria appears to be driven mainly by the negative effect of sulfate (presumably more due to its osmotic effect than to specific toxicity). Conductivity, widely acknowledged as a key factor influencing periphytic communities, particularly diatoms, is underscored by several studies (Soininen et al., 2004; Virtanen and Soininen, 2012; Stenger-Kovács et al., 2020). It mainly reflects the overall ionic concentrations and closely correlates with water pH, another important factor noted in this study, as besides these two main environmental gradients, pH was positively correlated with the total and cyanobacterial biomass estimates; and diatom biomass estimates negatively with water temperature (as well as ammonium that can be seen as a component of the above water chemistry gradient).

We used two statistical approaches (PERMANOVA and RDA) to identify the main environmental drivers of community variability in three data sets: diatoms as assessed by microscopy; diatoms assessed by amplicon sequencing; and all microalgal taxa by amplicon sequencing. Common to all PERMANOVA and RDA results for all three data sets was the identification of a strong role of conductivity in driving patterns of community variability in both diatoms and all microalgal taxa (Table 2, Figs 5 and 7), reflecting the main water chemistry gradient described. Both PERMANOVA and RDA highlighted a strong response of diatom and microalgae communities to pH and water temperature besides conductivity in the case of both the amplicon and the microscopy data sets.

In the case of the microalgal amplicon data, discharge, total phosphate concentrations showed significant explanatory power of community composition besides conductivity, water temperature and pH, indicating possible taxon-specific differences in ecological responses of different microalgal groups. Cyanobacteria, which were found to respond to some environmental gradient in terms of their pigment signatures, were not captured by our community data sets at all, leaving their specific responses open.

In addition, RDA showed that flow velocity was a significant factor for microalgae community composition. This is consistent with earlier research, several studies have reported that stream flow velocity is a major factor influencing the heterogeneity in the composition of microphytobenthic communities and biodiversity in streams (Boulêtreau et al., 2010; Townsend et al., 2012; Breuer et al., 2017). Benthic microalgae are directly exposed to increased flow velocity and material movement carried by water current. Furthermore, gas bubbles produced through metabolic processes like photosynthesis or respiration can further disrupt and upset the stability or integrity of the microphytobenthic communities under high flow velocity conditions (Boulêtreau et al., 2010). The microalgae communities tend to be highly specialized at high flow velocity (Soininen, 2005). For example, attached taxa (like *Achnantidium minutissimum*) can tolerate higher flow velocity better than biraphid taxa (like *Navicula lanceolata*), which live on the substrata. Flow velocity influences the availability of other resources, such as light, suspended particles and nutrients, as well as other abiotic factors (e.g. temperature) (Townsend et al., 2012; Breuer et al., 2017), thus having a significant impact on microalgae community composition.

Judged qualitatively by the spread of points in the RDA, as well as by the number of significant environmental correlations revealed, or also by the number of OTUs being almost an order of magnitude higher than the number of taxa observed microscopically, the amplicon data set provided a better resolution of the ecological signals in diatom communities than the microscopy data set (Zimmermann et al., 2015). Although neither taxa observed by microscopy nor OTUs can be expected to strictly depict “real” species, this might indicate that the higher richness captured by the amplicon method at least partially contains ecologically relevant information beyond that contained in the microscopy data. Since closely related species often have identical 18S rDNA sequences, it could be expected that even this signal could possibly be improved by the use of more variable markers (Zimmermann et al., 2014). By the same criteria, with slightly different main drivers revealed in the PERMANOVA, the ecological resolution of the microalgal amplicon data set seems to have been the highest. This might be expected simply due to the broader taxonomic coverage.

The main gradients explaining community variability in the RDA ordinations were highly similar across the three data sets (with most variation in depicted patterns corresponding to rotation and mirroring of highly similar configurations). Common to all three data sets, these ordinations differed from what might have been expected from the environmental PCA by showing a strong alignment between the effects of the above described main hydrological and water chemistry gradients, with water temperature and pH showing strong effects, close to orthogonal to this main axis of variability. This is in line with the PERMANOVA results, and could indicate that less prominent gradients (as identified by a PCA) can also have strong ecological effects. As to the alignment of the main hydrological and water chemistry axes of variability, it is possible that more extensive sampling could differentiate between their effects upon community composition more clearly, this aspect will need to be revisited in the future. These findings agree with other studies in which water physical and chemical variables were main drivers of changes in microalgae assemblages, often with shift from more sensitive species to more tolerant species composition (Soininen et al., 2004; Virtanen and Soininen, 2012; Teittinen et al., 2015; Breuer et al., 2017).

The community ordinations (RDAs) also delivered somewhat unexpected results with respect to the FAL1, KRE1, KRE2 group of samples. These appeared with a substantial offset from all other samples in the environmental PCA, indicating a large difference in terms of both nutrient and salt concentrations. Although they appeared in relatively close vicinity of each other also in the RDA plots, their outlier position compared to all other samples was less pronounced than in the environmental PCA. Also in this respect, one can observe a pattern of increasing resolution in the rank order of diatom microscopy; diatom amplicon; microalgal amplicon data set.

Our results are consistent with (Nguyen et al., 2023), who found that the macroinvertebrate communities in the Kinzig catchment followed an environmental gradient ranging from sites with good, moderate to poor water quality. For example, the majority of species in the macroinvertebrate communities found at good water quality sites were grazers that preferred clear water with coarser substrates, and oligotrophic nutrient content. On the other hand, species that were foragers or active filter feeders and had a preference for turbid water with finer substrates, brackish and eutrophic/mesotrophic conditions were more likely to be found in poor water quality sites (Nguyen et al., 2023).

Our IPS results from microscopy analysis showed poor to good ecological status, while amplicon analysis showed poor to high (very good) ecological status at the sampled sites throughout the catchment (Figures 5 and 7A). This could be explained by the discrepancies in community composition observed between the two methods. These discrepancies may be due to biases in both methods. For example, microscopy may ignore some barely visible taxa, while amplicon sequencing still miss many taxa in the PR2 reference library used in the present study (Zimmermann et al., 2014). Diatom ecological status indices such as the IPS are calculated on the notion that the ecological quality of a site is represented by the average ecological value of its taxa, weighted by their relative abundances (Stevenson et al., 2010). As a result, high abundance of a taxa strongly influences these indices, whereas low abundance of a taxa only slightly affects the final value (Tapolczai et al., 2024). For example, when our microscopic observations revealed that *Navicula lanceolata* was the most dominant taxa in most of the samples, amplicon analysis showed that the most abundant OTUs were associated with *Navicula radiosa*. And, the latter had a higher sensitivity to the IPS (S = 5.0) than the former (S = 3.8). The variations in water quality classes between the two methods may also be due to overestimation of the relative abundance of large species – for example, the *Achnanthidium minutissimum* case mentioned above. However, our results were consistent in the case of site KRE2, which had a poor ecological status as demonstrated by both microscopy and amplicon analysis. The substrate at this site was largely sediment and it is one of the three sites discriminated by the PCA, located at the lowest altitude, which followed a nutrients and salinity gradient.

### 4.2 Taxonomic inconsistencies between diatom microscopy and amplicon sequencing

Our results showed that microalgae communities characterized by different methods used in this study were influenced by various environmental factors, which agrees with previous studies (Virtanen and Soininen, 2012; Teittinen et al., 2015; Stenger-Kovács et al., 2020). Digital microscopy analysis revealed that *Navicula lanceolata* was the most dominant diatom taxon in the majority of our sampling sites, except KIN3 and KIN4. This taxon was reported to be the most dominant in an anthropogenically modified river in Poland, reaching over 80% of the total number of counted valve in spring (Noga et al., 2013). The authors attributed this dominance to the sampling season rather than anthropogenic influences. It was also identified as one of the taxa that had a higher proportion in agricultural land, with a higher content of nutrients and ions, such as in the Carpathian Basin (Stenger-Kovács et al., 2020). It occurred massively in slightly eutrophic boreal streams (Virtanen and Soininen, 2012). Due to its broad trophic preference for electrolyte-rich water and cooler temperatures, *N. lanceolata* grows rapidly during the winter and spring months (Noga et al., 2013; Schröder et al., 2015; Cantonati et al., 2017). This may explain why this species was highly represented in our samples collected in the first weeks of spring 2021. Therefore, the dominance of this taxon in our samples could also be related to the sampling season.

In contrast, amplicon sequencing analysis showed that *Navicula radiosa* was the most dominant taxon in most of our samples. We did not identify this taxon in microscopic analysis, instead we found *N. lanceolata* to be the most abundant taxon. Given the common taxonomic grouping of these two diatom species in the genus *Navicula*, they certainly have some similar morphological features that could lead to confusion in identification. However, *N. lanceolata* has obtusely rounded ends and a roundish central area (Cantonati et al., 2017). The observed discrepancies in our results are likely due to misidentification in the reference database, suggesting that the name *N. radiosa* may have been incorrectly assigned to OTUs belonging to *N. lanceolata* (Zimmermann et al., 2014). Other taxa detected in high abundances through molecular analysis in this study, include *Navicula gregaria and Surirella brebisonii*. These two taxa were also identified in light microscopy analysis. They grow abundantly in various environmental conditions, from mesotrophic to eutrophic waters. For example, *N. gregaria* has been reported as one of the indicators of nutrient enrichment in boreal alkaline and eutrophic basins (Virtanen and Soininen, 2012). We suspect that the presence of this taxon in large quantities in our samples may be related to nutrients and salt concenration in the rive catchment.

We found 1400 different diatom OTUs, about ten times the number of taxa we detected with morphological observations (142), demonstrating the broader taxonomic coverage of amplicon analysis. However, the reference sequences from the 18S-V9 region for many freshwater diatom taxa are still missing from the PR2 reference database used in the present study. This shows that a lot of work is still needed to complete the reference databases in this research area. Even the well-curated and updated diatom reference database as Diat.barcode (Rimet et al., 2019) is still incomplete, which can lead to both non-identification and misidentification of taxa (Zimmermann et al., 2014; Bailet et al., 2019). In addition, it is not clear whether the issues of cryptic and pseudocryptic species and infraspecific or intragenomic variation are covered by rDNA reference databases (Minerovic et al., 2020).

Our results showed that only a small number of diatom taxa could be identified through both morphological analysis and amplicon sequencing to the lowest taxonomic level. The reference database used for taxonomic assignment in our amplicon analysis is likely incomplete, which could explain the large differences in taxonomic composition between the two methods. Since diatoms are unicellular organisms, variables such as the number of genomes and the number of gene copies per genome influence the number of copies of a gene (Vasselon et al., 2018). This implies that gene copy number could be approximated by cell biovolume. From the smallest to the largest diatom species, there is variation in gene copy number that can have a major effect on metabarcoding quantification (Vasselon et al., 2018). This could explain why highly abundant small-size taxa observed in microscopy such as *Achnanthidium minutissimum* were not as dominant in amplicon analysis.

In conclusion, our study aimed to identify the key environmental variables that influence microalgal biomass and community composition in the Kinzig catchment. Our results showed that depth gradient slightly negatively correlated with total and green algal biomass. Sulfate showed a strong negative correlation with the total, cyanobacterial, and diatom biomass. Total and cyanobacterial biomass were positively associated with pH, while diatom biomass was negatively correlated with ammonium content and water temperature. PERMANOVA showed that conductivity, water temperature and pH were the most important factors influencing microalgae community composition, as observed in both microscopy and amplicon analysis. Together with these three variables, total phosphate in all microalgae OTU and water discharge in diatom (Bacillariophyta) OTU adatasets may suggest taxon-specific variations in the ecological responses of various microalgae groups. Our results highlighted the sensitivity of eukaryotic algae to physical and chemical variables, demonstrating their potential as good indicators of environmental gradients in the Kinzig catchment. These results suggest the complex relationship between multiple environmental factors and microalgal biomass and community composition. Further investigations, including collection of time series data, are required to fully understand the underlying biotic and abiotic factors driving these microalgal communities.

## Data availability

The datasets presented in this study can be found in the online repository Zenodo: https://doi.org/10.5281/zenodo.11061732

## Conflict of Interest

The authors declare that the research was conducted in the absence of any commercial or financial relationships that could be construed as a potential conflict of interest.

## Author Contributions

NASM: Data curation, Formal analysis, Investigation, Methodology, Software, Validation, Visualisation, Writing – original draft, Writing – review & editing. MD: Data curation, Formal analysis, Writing - review & editing. MK: Investigation, Methodology, Software, Writing: review & editing. DV: Data curation, Formal analysis, Writing: review and editing. DB: Investigation, Methodology, Software, Writing: review & editing. AMBC: Investigation, Methodology, Supervison, Formal analysis, Validation, Writing – review & editing. BB: Conceptualization, Funding acquisition, Supervision, Project administration, Writing – review & editing.

## Funding

This study was performed within the framework of the Collaborative Research Centre (CRC) 1439 RESIST (Multilevel Response to Stressor Increase and Decrease in Stream Ecosystems; www.sfb-resist.de) funded by the Deutsche Forschungsgemeinschaft (DFG, German Research Foundation; CRC 1439/1, project number: 426547801).

## Acknowledgements

We want to thank Marzena Spyra and all the students who helped with sample collection and in the laboratory with sample processing. The editor and two anonymous reviewers are thanked for their comments that helped improve this manuscript. We acknowledge support by the Open Access Publication Fund of the University of Duisburg-Essen.

## Supplementary materials

**Supplementary Table 1** Physical and chemical parameters analysed in this study

**Supplementary Table 2** List of taxa idedntified by microscopy (_MIC) or amplicon sequencing (_DNA) with their 4 letters OMINIDIA codes

## Notes

https://doi.org/10.5281/zenodo.11061732

### Competing Interest Statement

The authors have declared no competing interest.

